# *AdamTS-B* protease is required for morphogenesis of the *Drosophila* respiratory system

**DOI:** 10.64898/2026.08.19.745706

**Authors:** Jana Schulze, Uwe Töpfer

## Abstract

Epithelial tube morphogenesis is critical for the function of many organs. Basement membranes underlie epithelia and their remodeling is a key step to reach the correct size and shape. Key regulators that mediate basement membrane remodeling for tube elongation and branching remain largely unknown. We analyze the expression and function of *AdamTS-B*, a matrix metalloprotease, in the respiratory system of *Drosophila*. Here we show, that *AdamTS-B* is expressed early in tracheal development during placode formation. We generated a mutant line of *AdamTS-B*, which is lethal. Analysis of trachea morphogenesis in this *AdamTS-B* mutant reveal a function in tube elongation and cell migration. Our results suggest that *AdamTS-B* control BM remodeling required for organ shape.

## Introduction

The basement membrane (BM) is a thin extracellular matrix that underlies epithelia and surrounds other tissues (Jayadev and Sherwood, 2017; Töpfer, 2023). These extracellular matrices provide cells with biochemical and mechanical information and serve as a scaffold for cell adhesion, act as barrier and influence cell signaling as well as cell behavior, such as proliferation, migration and differentiation (Page-McCaw and Ferrell, 2025; Sherwood, 2021; Yurchenco, 2011). They are made primarily of collagen IV, laminin, nidogen, and heparan sulfate proteoglycans such as perlecan, which self-assemble in sheet-like structures (Hohenester and and Yurchenco, 2013).

AdamTS proteases (A Disintegrin and Metalloproteinase with Thrombospondin motifs) are a family of zinc-dependent enzymes that play a central role in the formation and breakdown of tissue in the human body. They play a key role in the formation, homeostasis and remodeling of the extracellular matrix. The substrate spectrum of AdamTS proteases can range from individual to entire families of extracellular matrix proteins (Taye et al., 2023). To date, 19 different variants of these enzymes are known in humans (Kelwick et al., 2015). This matrix proteases have a diverse role in tissue morphogenesis and human pathologies. For example, dysregulations of AdamTS proteases are associated with different diseases in humans, like arthritis (Lin and Liu, 2010), cancer (Kumar et al., 2012; Rocks et al., 2008) or atherosclerosis (Bauer et al., 2015; Stanton et al., 2011). Using genetic model organisms, it has been shown that AdamTS proteases are important for cell migration and organ development in various tissues (Agarwal et al., 2022; Gonsior and Ismat, 2019; Ismat et al., 2013; Ku and Bilder, 2025; Lhamo and Ismat, 2015; Töpfer et al., 2024). *AdamTS-B* is one of four *Drosophila* AdamTS genes (Öztürk-Çolak et al., 2024). Knockdown of *AdamTS-B* in the *Drosophila* wing disc leads to a disruption in wing vein formation, which has been associated with a previously identified role in the regulation of the EGFR signaling pathway (Butchar et al., 2012; Pham et al., 2018; Schulze and Töpfer, 2025).

The development of the tracheal system in the *Drosophila* embryo is a powerful model for studying epithelial morphogenesis and organogenesis. Here, genetic and mechanical cues work together to transform simple epithelial primordia into a highly branched, functional respiratory organ. From around stage 10 of embryogenesis, clusters of ectodermal cells invaginate to form tracheal placodes. These placodes then undergo coordinated cell migration and branching morphogenesis to generate a complex tubular network that delivers oxygen throughout the organism. This process is guided by finely tuned genetic programs and signaling pathways, including Fibroblast growth factor (FGF) signaling, which directs outgrowth and branching and Wingless and Decapentaplegic (Dpp) pathways determine the fate of the trachea (Llimargas, 2000), EGFR signaling controlling tube elongation by regulation of intracellular trafficking as well as cell migration (Llimargas and Casanova, 1999; Olivares-Castiñeira and Llimargas, 2017) and Integrin-BM interactions stabilize the structure (Levi et al., 2006; Sánchez-Cisneros et al., 2025). In the final stages of embryonic development, remodeling of the apical matrix, controlled through SRC signaling, ensures proper cell shape changes which are required for tube elongation (Förster and Luschnig, 2012).

Here we identified the matrix protease AdamTS-B as an important regulator of trachea morphogenesis in the *Drosophila* embryo by modulating the cell migration of ganglionic and visceral branches as well as mediating the shape of the dorsal trunk. We show the unique expression of *AdamTS-B* during embryonic development starting form differentiated placodes of trachea precursor. We generated a *AdamTS-B* mutant allele using CRISPR/Cas and revealed lethality of homozygous *AdamTS-B* mutant flies. Together, these results identify AdamTS-B as a miscellaneous modulator of tracheal morphogenesis.

## Results

### *AdamTS-B* expression in trachea

The embryonic development of *Drosophila* is a rapid morphogenetic process in which the larval organs are formed within 24 hours. During this process, the developing BM undergoes a remodeling that plays a crucial role in the proper formation of these organs (Martin et al., 1999; Urbano et al., 2009; Wolfstetter and Holz, 2012; Wolfstetter et al., 2019; Yarnitzky and Volk, 1995). To determine whether the AdamTS protease *AdamTS-B* is expressed during embryonic development, we used a genetic tool that extremely sensitively reflects induction of expression activity. CRIMIC (CRISPR-Mediated Integration Cassette) lines contain a T2A-GAL4 cassette to drive GAL4 expression matching the host gene’s pattern, a series of genetic tools that is called Trojan-Gal4 (Kanca et al., 2019; Lee et al., 2018). We first detected reporter gene expression in stage 11 embryos, the timepoint when ectodermal tracheal placodes have been formed (Fig. 1A, B and B’). This expression is detectable in the entire respiratory organ until the end of embryonic development (Fig. 1C-F’). We did not find any other tissue showing reporter gene expression, which indicates that *AdamTS-B* is solely expressed in trachea cells. The early expression of *AdamTS-B* may indicate a role in the remodeling of the tracheal BM during development.

**Figure 1:**
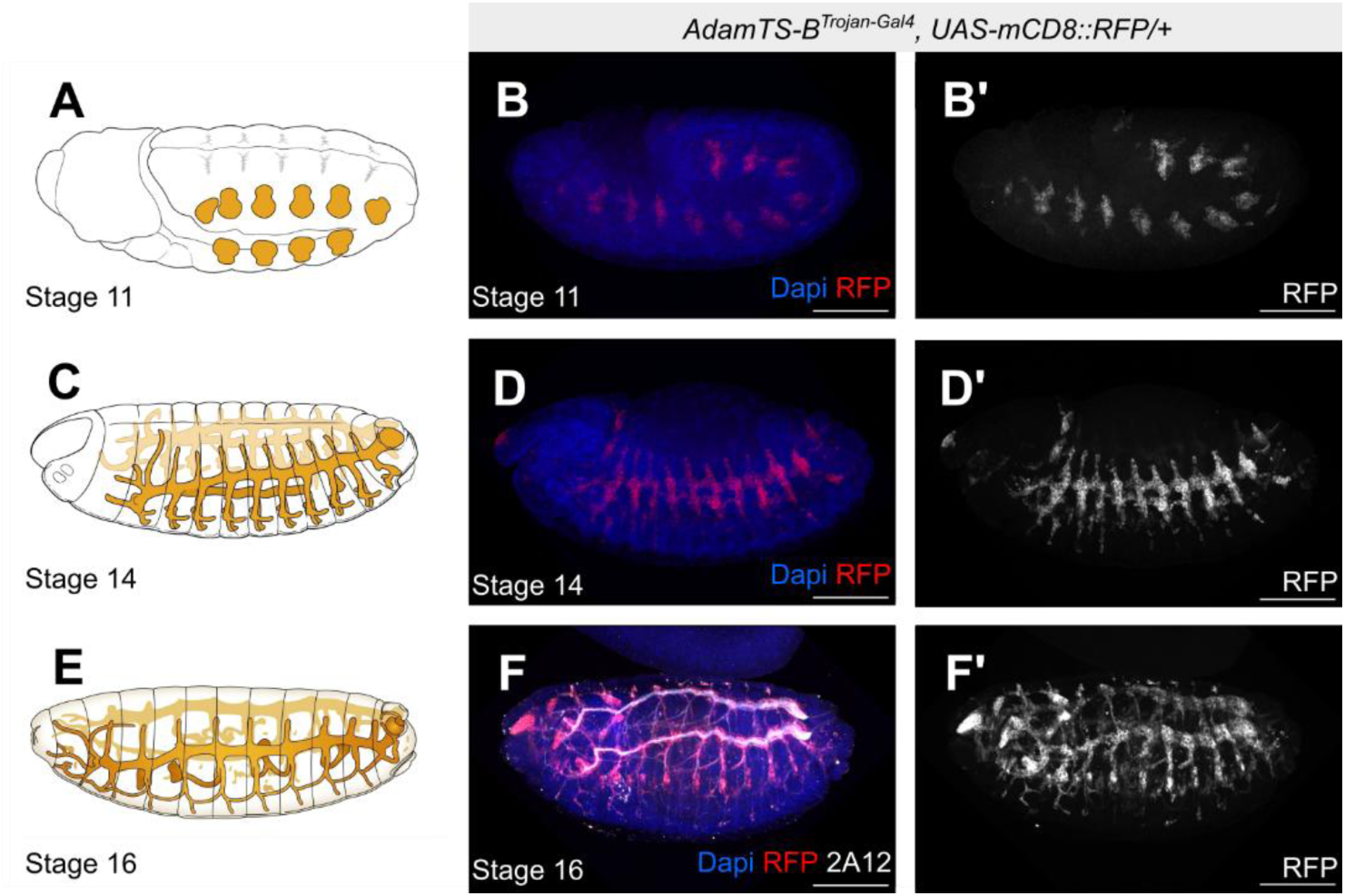
*AdamTS-B* Trojan insertion reveal expression in tracheal embryonic cells. **(A, C, E)** Schematic illustration of trachea (orange) development in the embryo. Adopted after (Hartenstein, 1993). **(B, B’, D, D’, F, F’)** Embryos with the genotype *AdamTS-B^Trojan-Gal,^ UAS-mCD8::RFP/+* in stage 11 (B and B’), stage 14 (D and D’) and stage 16 (F and F’) stained with Dapi (blue), RFP (red) and anti-2A12 (white) for the apical matrix of trachea. Scale bars = 100 µm.

### AdamTS-B^KO^ is lethal

The BM has previously shown to affect various aspects of trachea development. Mutants for laminins show phenotypes regarding the shape or gaps of the of the dorsal trunk or disturbed cell migration (Klußmann-Fricke et al., 2022; Martin et al., 1999; Urbano et al., 2009; Wolfstetter and Holz, 2012). To determine whether *AdamTS-B* plays a significant role in the respiratory system, we generated an *AdamTS-B* mutant using non-homologous end-joining repair with CRISPR/Cas9, henceforth referred to as *AdamTS-B^ko^* (Fig. 2A). Next, we wanted to test if these mutant leads to lethality in any stage of *Drosophila* development. We used a Balancer chromatid with a fluorescent marker which is detectable in larvae, pupae as well as in adult flies. The expected ratio of 33% fell already below this level in early L1-stage larvae and continued to decline through the third larval stage, while no homozygous *AdamTS-B* mutant individuals were found in either pupae or adult flies (Fig. 2B). Thus, *AdamTS-B^ko^* mutant animals die over larval development.

**Figure 2:**
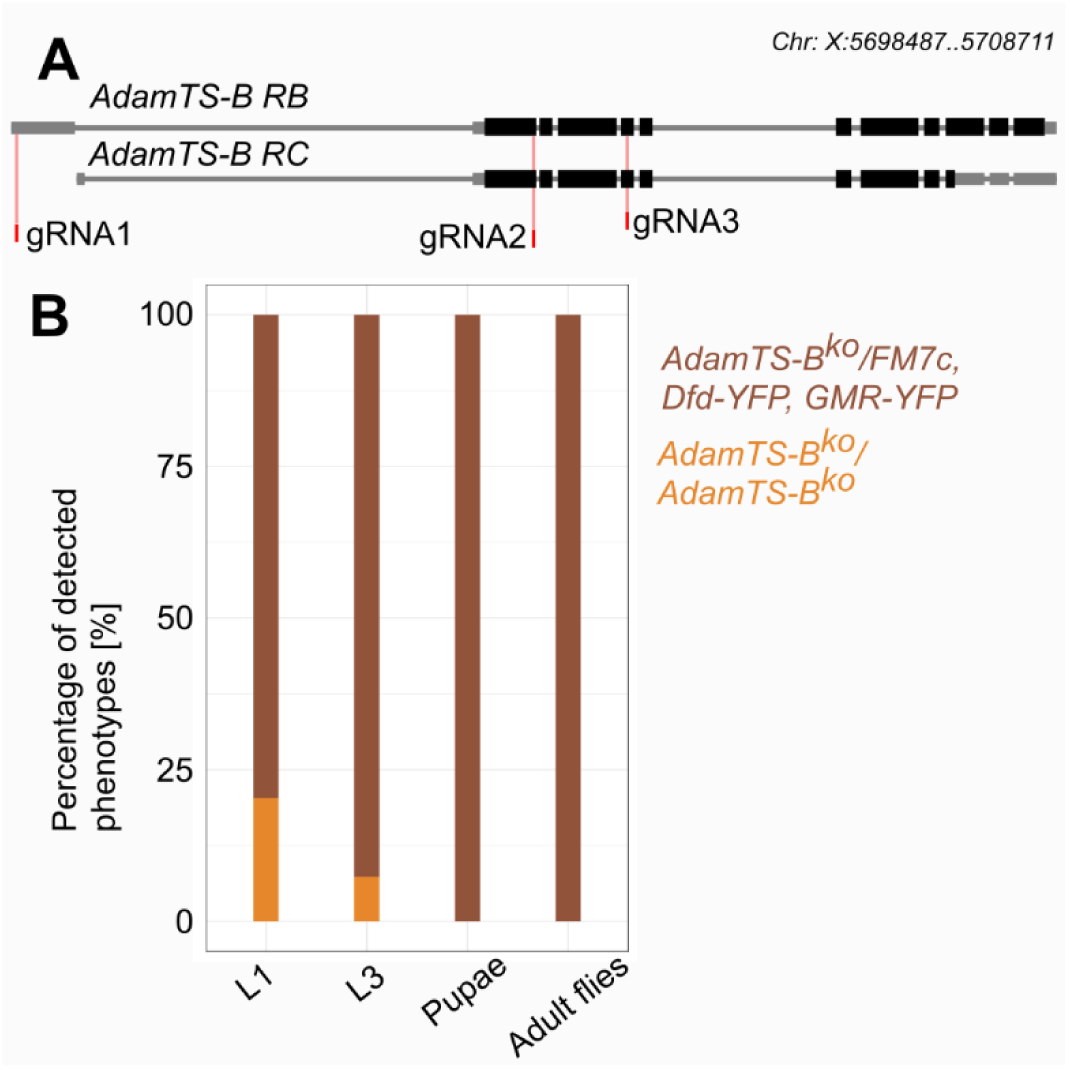
*AdamTS-B* mutants are larval lethal. **(A)** Scheme of *AdamTS-B* genomic region with two known isoforms (RB and RC) and loci of gRNA targets. **(B)** Percentage of phenotypes detected in inter-se crossings from parental flies with the *AdamTS-B^ko^/FM7c, DFD-YFP, GMR-YFP* genotype in distinct stages of development. *n* > 300 per stage and genotype.

### AdamTS-B regulates proper trachea morphogenesis

To understand whether the embryonic expression of *AdamTS-B is* associated with the lethality of the mutant, we studied the morphogenesis of the tracheae. We used an anti-GASP (2A12) antibody that stains the apical matrix of the trachea lumen between stages 14-17 in control (Fig. 3A-D) and *AdamTS-B* mutant embryos (Fig. 3E-H) (Manning and Krasnow, 1993).

**Figure 3:**
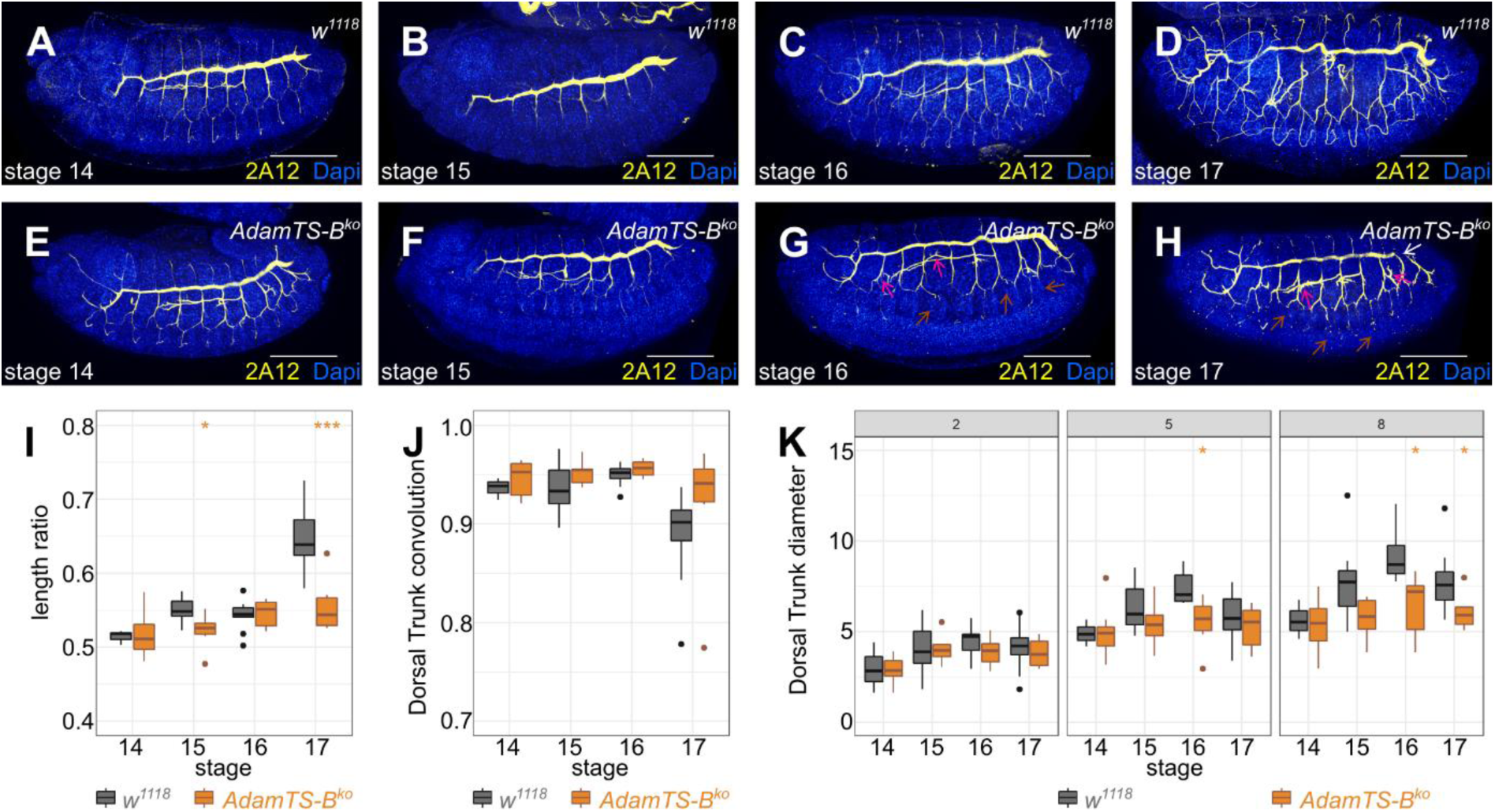
AdamTS-B is required for proper trachea cell migration and organ formation. **(A-H)** Staining of laterally aligned embryos of *w^1118^* (A-D) and *AdamTS-B^ko^*(E-H) was performed for the nuclei with Dapi (blue) and 2A12 for the apical matrix (yellow) in stages 14 to 17. Arrows are pointing towards gaps within the DT (white), multiple branching of the VB (magenta) and shorter branches of the LT (brown). **(I-K)** Quantifications of the length ratio (I), the DT convolution (J) and the diameter of the DT at positions 2, 5 and 8 (K) for the indicated genotypes and stages. *n* > 12 embryos per genotype. Significances were estimated using the two-tailed students t-test. Significance levels: * = p < 0.05, ** = p < 0.01, *** = p < 0.001, scalebar: 100 μm.

First, we studied the branching of the migrating tracheal cells, which generate a stereotypical pattern of branches that are connected to the dorsal trunk (DT) including the transversal connective (TC), lateral trunks (LT), dorsal branches (DB), visceral branches (VB) and the ganglionic branch (GB) (Fig. 3A-D, Sup. Fig. 1). In stages 16 and 17, we found an abnormal multiple branching of the VB in *AdamTS-B* mutants (Fig. 3G, H; magenta arrows). Additionally, LT and GB branches seemed to be shorter in these stages compared to the control (Fig. 3G, H brown arrows). Second, we quantified the dimensions of the DT. To examine this, the DT was analyzed using three following parameters: length ratio, convolution, and diameter (Sup. Fig. 2). The last parameter is further divided into three distinct positions as the diameter of the DT ranges also in the wild type and gets thicker from anterior to posterior (Hayashi and Kondo, 2018). Therefore, one position near the anterior of the DT (position 2,) one in the middle (position 5) and one near the posterior of the DT (position 8) were measured (Sup. Fig. 2). In stages 14 and 16, the length ratio, as well as the convolution, did not show a significant difference between the control and the loss of *AdamTS-B* (Fig. 3A, C, E, G, I, J). Despite that, the DT of *AdamTS-B^ko^* was significantly shorter than in the controls in stages 15 and 17. In contrast, the convolution showed no significant differences in these stages (Fig. 3B, D, F, H, I, J). The effects related to the diameter of the DT in *AdamTS-B^ko^* were observed to be position-dependent. At the second position no significant difference could be detected in all stages (Fig. 3K, position 2). At position 5 and 8, diameter increases from stage 4-16 and drops again in 17. This thickening of the DT between stages 14 and 16 is less pronounced in the mutant, leading to significant differences in stage 16 (Fig. 3K). In stage 17, however, a significant difference can only be observed in position 8 (Fig. 3K). Additionally, gaps within the DT were observed in all stages of *AdamTS-B^ko^*(exemplary seen in Fig. 3H white arrow). Overall, these results show that *AdamTS-B* has an impact on the development of the tracheal system mainly in stage 17.

### *AdamTS-B* overexpression disturbs trachea branching and tube growth

Loss of AdamTS-B leads to altered DT properties regarding the length ratio and the diameter and disturbed branching (Fig. 4). This raises the question if an increased expression shows the opposite or the same phenotype as the loss of *AdamTS-B* expression. Therefore, we overexpressed *AdamTS-B* using the Gal4/UAS system with a trachea-specific GAL4 driver line (*btl-Gal4*).

**Figure 4:**
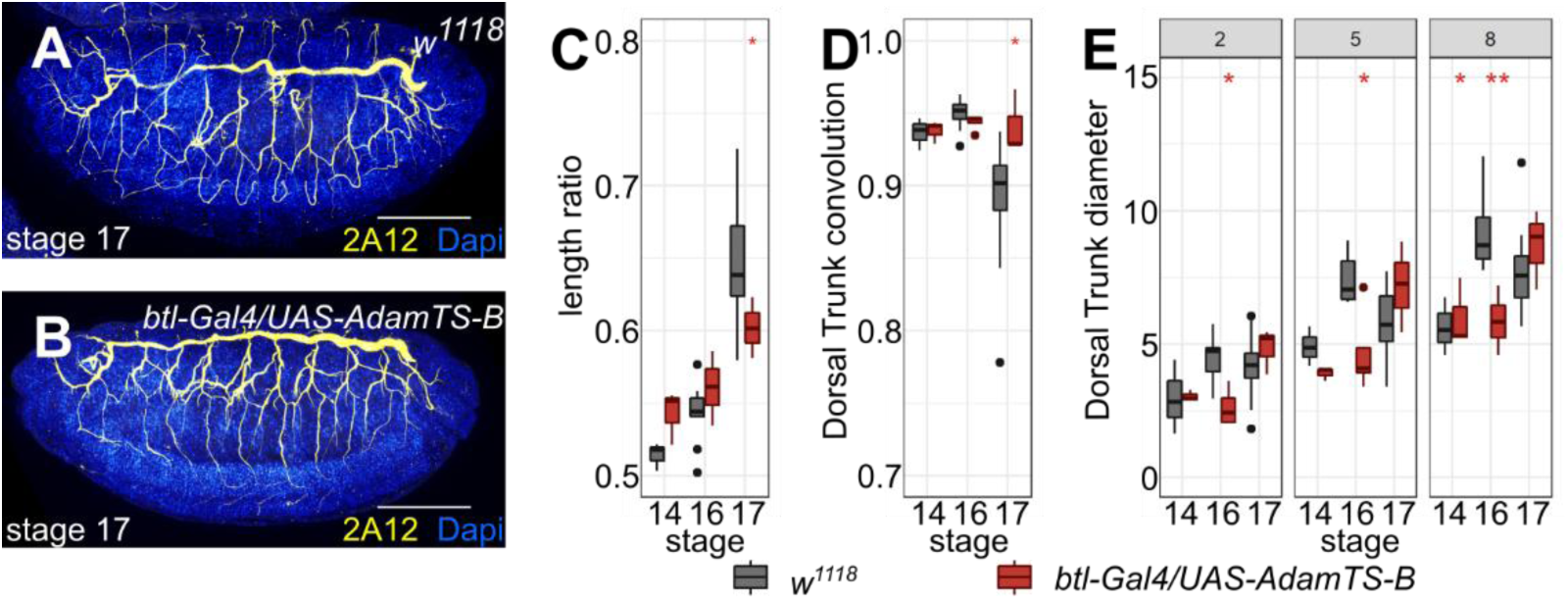
Overexpression of *AdamTS-B* disturbs epithelial tube elongation and cell migration. **(A and B)** Stage 17 Embryos of the control (A) and overexpression of *AdamTS-B* (B) are stained for the nuclei with Dapi (blue) and the tracheal lumen with 2A12 (yellow). **(C-E)** Quantification of the length ratio (C), the DT convolution (D) and the DT diameter at positions 2, 5 and 8 (E) were performed in stages 14, 16 and 17 for the indicated genotypes. *n* > 3 embryos per genotype. Significances were estimated using the two-tailed students t-test. Significance levels: * = p < 0.05, ** = p < 0.01, *** = p < 0.001, scalebar: 100 μm

In stage 17, the length ratio was significantly smaller in *AdamTS-B* overexpression than in controls (Fig. 5A-C). In concomitant with the lower length ratio, the DT was straighter in stage 17 embryos overexpressing *AdamTS-B* (Fig4. A, B, D). Furthermore, wildtype embryos and those overexpressing *AdamTS-B* exhibited distinct developmental trends in DT diameter at all positions. While the wildtype embryos first increased in diameter, followed by a decrease, the overexpression of the protease led to a steady diameter in stages 14 and 16, followed by an increase (see Fig. 5E). These discrepancies resulted in significant differences in stage 16 at all measured positions. A significantly different diameter was also found in stage 14 at position 8. (Fig. 4E). These results indicate the complex role of AdamTS-B in tracheal development and demonstrate the requirement for AdamTS-B to maintain the integrity of the DT.

## Discussion

Branching morphogenesis is one of the most multifaceted cell biological process to allow organs and tissues to form into highly complex structures that are required to fulfill their physiological functions (Goodwin and Nelson, 2020). On the cellular level branching morphogenesis is the results from the remodeling of epithelia that undergo directed cell proliferation, migration and cell shape changes (Affolter et al., 2003). In order to allow this epithelial remodeling, the components of the BM must also be remodeled (Loganathan et al., 2020). In the *Drosophila* tracheal system, proper BM integrity is essential for the development of the correct shape of the trachea (Klußmann-Fricke et al., 2022). The work described here, uncovers the function of a previously unknown regulator of trachea morphogenesis. Specifically, we show that *AdamTS-B* is expressed in ectodermal tracheal placodes, and that *AdamTS-B* mutant embryos die during larval development. In particular *AdamTS-B* knockout embryos have defects in branching morphogenesis as well as in tube elongation. Finally, our data shows that overexpression of *AdamTS-B* in the tracheae also leads to defects in tracheal development, but that these defects differ from those caused by the loss of the gene.

The increase in lethality during larval development that we observed can be explained by the tracheal defects during embryogenesis. Shortly before hatching, a *Drosophila* embryo drains the fluid from its respiratory tubes and quickly fills the tracheal network with gas. This vital transition transforms the tubes from a non-functional, fluid-filled state into an open respiratory system (Tsarouhas et al., 2007). The decrease in survival mainly occurs during larval development, when the respiratory organ is needed for survival.

*AdamTS-B* have been shown to act as a negative regulator of EGF signaling (Butchar et al., 2012). The phenotypes, we observed in our *AdamTS-B* mutant line in trachea development support the idea that AdamTS-B might also play a similar role in regulating EGFR signaling in the respiratory system. Overexpression of constitutive active from of EGFR leads to a shorter DT in stage 16 embryos (Olivares-Castiñeira and Llimargas, 2017). These results are consistent with our data, in which the loss of AdamTS-B also leads to shorter DT, supporting the idea of a negative regulator. Conversely, however, the overexpression of *AdamTS-B* in our experiment also results in a shorter DT. It has previously been shown, however, that the expression of a constitutively active EGFR leads to a much longer DT (Olivares-Castiñeira and Llimargas, 2017), which thus contradicts the hypothesis that *AdamTS-B* acts as a negative regulator. The molecular mechanism by which AdmaTS-B appears to cause BM and EGFR signaling to influence each other is therefore somewhat more complex.

In parallel on the completion of this manuscript, Thuringer and Steinmetz et al. published work that also examined the role of *AdamTS-B* on trachea morphogenesis (Thuringer et al., 2026). In particular, the analysis of gene expression in early tracheal precursors, which employed *in situ* hybridization rather than the Trojan tool we used, and the study of defects in branching in both a mutant context and in the case of *AdamTS-B* overexpression complement each other.

The expression and function of *AdamTS-B* in the *Drosophila* respiratory system raises the question if there might be a conserved function of AdamTS proteases between flies and human. *AdamTS-B* shows the closest relation to the human *AdamTS7* and *AdamTS12* (Sup. Fig. 3). Interestingly, *AdamTS12* is associated with asthma and bronchial hyperresponsiveness (Kurz et al., 2006; Paulissen et al., 2012). Thus, revealing functions of *AdamTS-B* in tracheal development could get further insights into the role of *AdamTS12* in lung development. Moreover, the tracheal system of *Drosophila* is a good model system for identifying novel mechanisms of chronic obstructive pulmonary disease (Scholl et al., 2021).

In this study we found a matrix protease that is required for the development of the *Drosophila* respiratory system. The exact mechanism by which AdamTS-B acts at the molecular level is still unclear, as is the precise manner in which the described defects occur. We propose a model where AdamTS-B is required to remodel the BM in a fashion that its changes the capacity to control morphogen distribution, which are required for branching morphogenesis and tube elongation (Hayashi and Kondo, 2018; Olivares-Castiñeira and Llimargas, 2017). The phenotypes and overexpression of *AdamTS-B*, as well as the previously demonstrated interdependencies between AdamTS-B and the EGFR signaling pathway, also suggest a role for this protease in the regulation of this pathway, which could be a common function of this group of AdamTS proteases also in vertebrates.

## Material and Methods

### Drosophila stocks

The fly stocks used were *w^1118^* (BDSC, 3605) [Bloomington Drosophila Stock Center (BDSC), *AdamTS-B^CR01623-TG4.2^* (BDSC, 86472), *FM7c, Dfd-GMR-YFP* (BDSC, 23229), *UAS-mCD8::RFP* (BDSC, 32219), *btl-Gal4* (BDSC, 8807), *nos-Cas9* (BDSC, 78781) and *UAS-*

*AdamTS-B* (Pham et al., 2018). Flies were kept at 25°C on standard food. Crosses were raised at 29°C.

### Immunohistochemistry

Antibody staining of *Drosophila* embryos was essentially performed as described in (Töpfer et al., 2019). The following primary antibodies were used: mouse anti-GASP (DSHB, 2A12, 1:5) and rat anti-RFP (Chromotek, 5F8, 1:1000). The secondary antibodies used were goat anti-mouse Cy3 (Jackson ImmunoResearch, 115-165-003, 1:200) and goat anti rat (Vector Laboratories, 1:500). DNA was stained with DAPI. Images were acquired using a Zeiss Axio Observer Z1 with Apotome. Embryos were embedded in Mowiol mounting medium with 2.5% DABCO.

### Generation of *AdamTS^KO^* line

The AdamTS-B mutant fly line were generated via CRISPR-Cas9 non homologues endpoint repair method. gRNAs were designed with E-CRISP (Heigwer et al., 2014) and cloned in pCFD5 (Addgene, 73914)(Port and Bullock, 2016). A mixture of all three gRNA vectors were injected in *Drosophila* embryos with nos-Cas9 genotype in a concentration of 100 ng/μl. The following primer to generate gRNA vectors were used: gRNA1 forward TGCAGCGGCTCTCT GTGGGAAGCG, gRNA1 reverse AAACCGCTTCCCACAGAGAGCCGC, gRNA2 forward TGCAGTGGCCATC TCCACATGCGC, gRNA2 reverse AAACGCGCATGTGGAGATGGCCA C, gRNA3 forward TG CAGGATCCGCATCACCATGACG, gRNA3 reverse AAACCGTCATGG TGATGCGGATCC.

### Analysis of dorsal trunk shape

All pictures were taken as Z stacks (distance: 1.24 µm) using the Observer.Z1 fluorescence microscope (air objective: 20x) with the Apotome.2. The analyzed pictures were processed to a maximum Z projections in Fiji (Schindelin et al., 2012). The DT was analyzed based on three different parameters. The length ratio was estimated by dividing the embryo’s length by the DT length between the first and ninth connecting points to the TC. The same length was used to calculate the convolution. The other parameter used was the ferret of this line (shortest possible distance between starting and ending points). The convolution was calculated by dividing the ferret by the length of the same line. The final parameter was the diameter, measured at three points: the second branching point of the DT to the TC in thorax segment 3 (position 2), the fifth branching point between the DT and the TC in abdominal segment 3 (position 5), and the eighth branching point in abdominal segment 6 (position 8). All measurements were performed using Fiji.

### Phylogenetic and protein domain analysis

The Protein domains were predicted with SMART (Letunic and Bork, 2026) and the phylogenetic tree was generated using HMMER (Rajković et al., 2026).

### Quantification and statistical analysis

Statistical significance was calculated using a two-sided students t-test in R (R Core Team, 2021).

## Acknowledgements

We thank Afshan Ismat for providing fly stocks. We are most grateful to Christian Dahmann for access to the microscope and infrastructure and Jinhee Ryu for the help with injection of AdamTS-B gRNAs. Stocks obtained from the BDSC (NIH P40OD018537) were used in this study. Antibodies were obtained from the Developmental Studies Hybridoma Bank, created by the NICHD of the NIH and maintained at The University of Iowa.

## Footnotes

### Author contributions

Conceptualization: U.T.; Data curation: J.S., U.T.; Formal analysis: J.S., U.T.; Investigation: J.S., U.T.; Methodology: J.S., U.T.; Project administration: U.T.; Resources: U.T.; Supervision: U.T; Validation: J.S., U.T.; Visualization: J.S., U.T; Writing – original draft: J.S., U.T.; Writing – review & editing: J.S., U.T.

### Data and resource availability

Data reported in this paper will be shared by the lead contact upon request.

## Supplementary Figures

**Sup. Figure 1:**
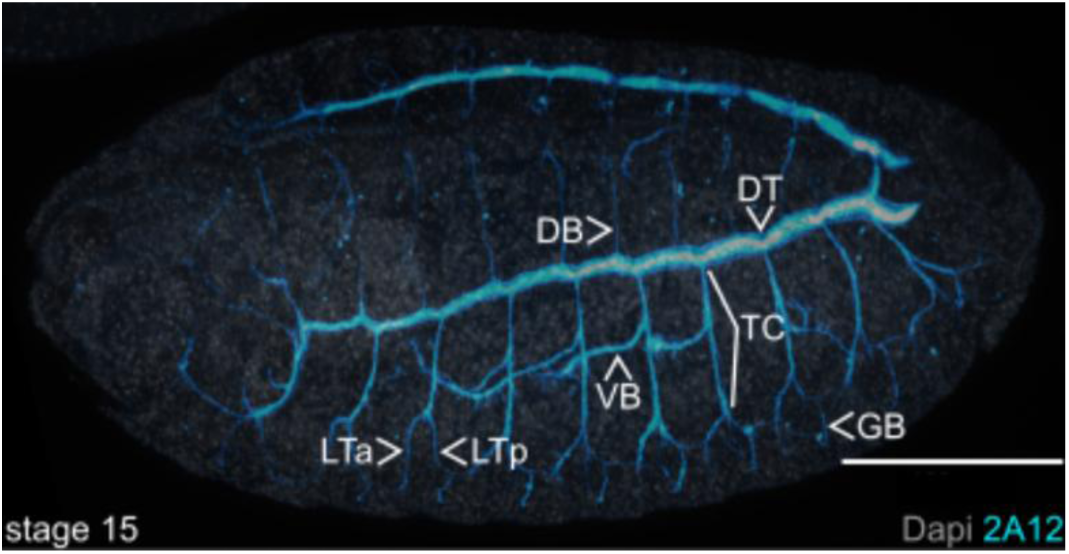
The tracheal system of *Drosophila* is composed of different specific branches. Shown is a basolateral aligned embryo at stage 15 stained for the nuclei (Dapi, grey) and the trachea (2A12, cyan). The dorsal trunk (DT) connects all branches. The transversal connective (TC) is the connection between the DT and the lateral trunks (LT) which can be divided into the LT anterior (LTa) and LT posterior (LTp). The ganglionic branch (GB) is connected to the LT and the visceral branch (VB) is a branch of the TC. Scalebar: 100 μm.

**Sup. Figure 2:**
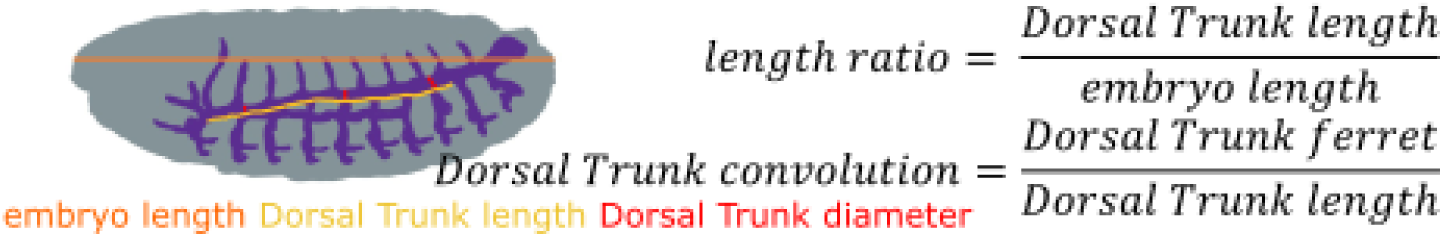
Schematic of the DT measurements. The length and the ferret of the DT (yellow) as well as the length of the embryo (orange) were measured. The diameter of the DT (red) was measured in the thorax segment 3 (position 2), the abdominal segment 3 (position 5) and the thorax segment 6 (position 8). Length ratio and DT convolution were estimated as indicated.

**Sup. Figure 3:**
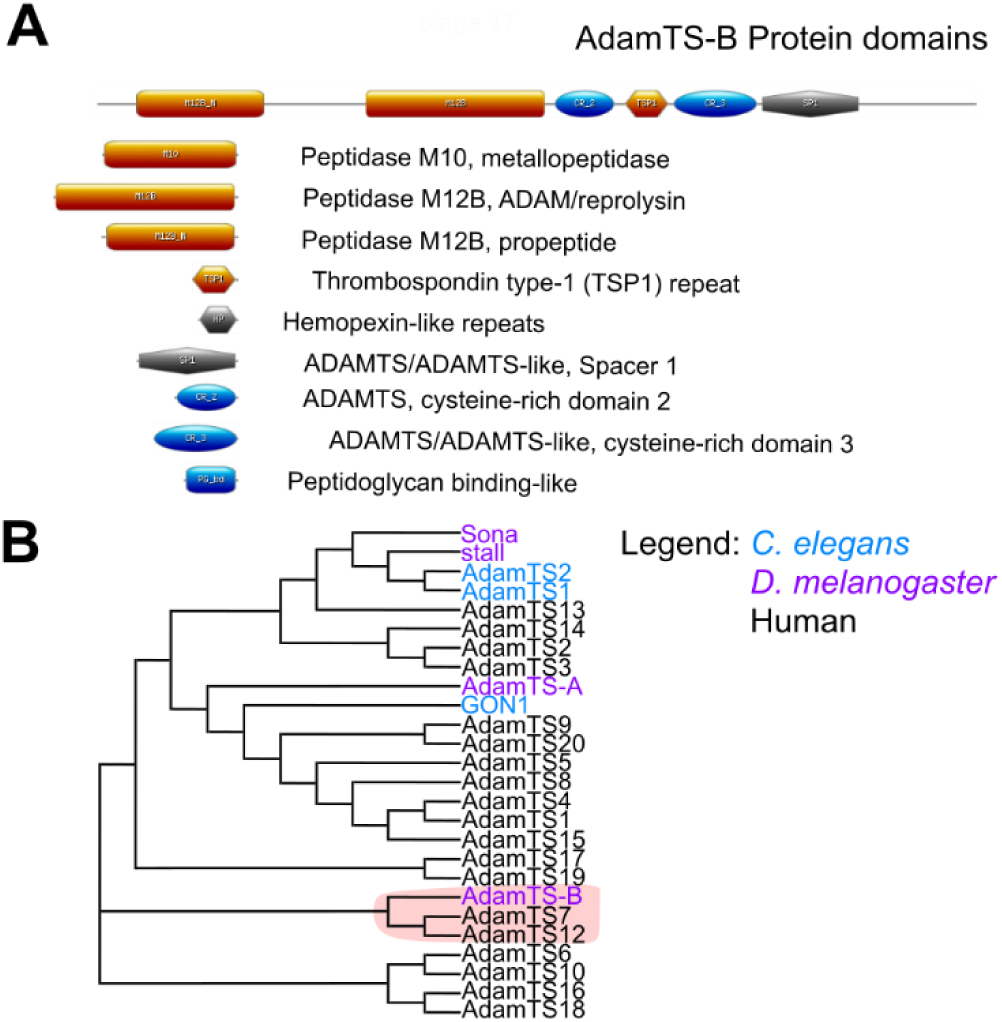
*AdamTS-B* is an ortholog of human *AdamTS7* and *AdamTS12*. **(A)** Scheme of AdamTS-B protein domains. (B) Phylogenetic tree of AdamTS proteases form human (black), *C. elegans* (blue) and *Drosophila* (magenta).

## References

Affolter, M., Bellusci, S., Itoh, N., Shilo, B., Thiery, J.-P. and Werb, Z. (2003). Tube or Not Tube: Remodeling Epithelial Tissues by Branching Morphogenesis. Developmental Cell 4, 11–18.

Agarwal, P., Shemesh, T. and Zaidel-Bar, R. (2022). Directed cell invasion and asymmetric adhesion drive tissue elongation and turning in *C. elegans* gonad morphogenesis. Developmental Cell 57, 2111–2126.e6.

Bauer, R. C., Tohyama, J., Cui, J., Cheng, L., Yang, J., Zhang, X., Ou, K., Paschos, G. K., Zheng, X. L., Parmacek, M. S., et al. (2015). Knockout of Adamts7, a Novel Coronary Artery Disease Locus in Humans, Reduces Atherosclerosis in Mice. Circulation 131, 1202–1213.

Butchar, J. P., Cain, D., Manivannan, S. N., McCue, A. D., Bonanno, L., Halula, S., Truesdell, S., Austin, C. L., Jacobsen, T. L. and Simcox, A. (2012). New Negative Feedback Regulators of Egfr Signaling in Drosophila. Genetics 191, 1213–1226.

Förster, D. and Luschnig, S. (2012). Src42A-dependent polarized cell shape changes mediate epithelial tube elongation in Drosophila. Nat Cell Biol 14, 526–534.

Gonsior, M. and Ismat, A. (2019). sli is required for proper morphology and migration of sensory neurons in the Drosophila PNS. Neural Dev 14, 10.

Goodwin, K. and Nelson, C. M. (2020). Branching morphogenesis. Development 147, dev184499.

Hartenstein, V. (1993). Atlas of Drosophila Development. Cold Spring Harbor Laboratory Press.

Hayashi, S. and Kondo, T. (2018). Development and Function of the Drosophila Tracheal System. Genetics 209, 367–380.

Heigwer, F., Kerr, G. and Boutros, M. (2014). E-CRISP: fast CRISPR target site identification. Nat Methods 11, 122–123.

Hohenester, E. and and Yurchenco, P. D. (2013). Laminins in basement membrane assembly. Cell Adhesion & Migration 7, 56–63.

Ismat, A., Cheshire, A. M. and Andrew, D. J. (2013). The secreted AdamTS-A metalloprotease is required for collective cell migration. Development 140, 1981– 1993.

Jayadev, R. and Sherwood, D. R. (2017). Basement membranes. Current Biology 27, R207–R211.

Kanca, O., Zirin, J., Garcia-Marques, J., Knight, S. M., Yang-Zhou, D., Amador, G., Chung, H., Zuo, Z., Ma, L., He, Y., et al. (2019). An efficient CRISPR-based strategy to insert small and large fragments of DNA using short homology arms. eLife 8, e51539.

Kelwick, R., Desanlis, I., Wheeler, G. N. and Edwards, D. R. (2015). The ADAMTS (A Disintegrin and Metalloproteinase with Thrombospondin motifs) family. Genome Biol 16, 113.

Klußmann-Fricke, B.-J., Martín-Bermudo, M. D. and Llimargas, M. (2022). The basement membrane controls size and integrity of the Drosophila tracheal tubes. Cell Reports 39,.

Ku, H.-Y. and Bilder, D. (2025). Basement membrane patterning by spatial deployment of a secretion-regulating protease. Proceedings of the National Academy of Sciences 122, e2412161122.

Kumar, S., Rao, N. and Ge, R. (2012). Emerging Roles of ADAMTSs in Angiogenesis and Cancer. Cancers 4, 1252–1299.

Kurz, T., Hoffjan, S., Hayes, M. G., Schneider, D., Nicolae, R., Heinzmann, A., Jerkic, S. P., Parry, R., Cox, N. J., Deichmann, K. A., et al. (2006). Fine mapping and positional candidate studies on chromosome 5p13 identify multiple asthma susceptibility loci. Journal of Allergy and Clinical Immunology 118, 396–402.

Lee, P.-T., Zirin, J., Kanca, O., Lin, W.-W., Schulze, K. L., Li-Kroeger, D., Tao, R., Devereaux, C., Hu, Y., Chung, V., et al. (2018). A gene-specific T2A-GAL4 library for Drosophila. eLife 7, e35574.

Letunic, I. and Bork, P. (2026). SMART v10: three decades of the protein domain annotation resource. Nucleic Acids Research 54, D499–D503.

Levi, B. P., Ghabrial, A. S. and Krasnow, M. A. (2006). Drosophila talin and integrin genes are required for maintenance of tracheal terminal branches and luminal organization. Development 133, 2383–2393.

Lhamo, T. and Ismat, A. (2015). The extracellular protease *stl* functions to inhibit migration of v’ch1 sensory neuron during *Drosophila* embryogenesis. Mechanisms of Development 137, 1–10.

Lin, E. A. and Liu, C.-J. (2010). The role of ADAMTSs in arthritis. Protein Cell 1, 33–47.

Llimargas, M. (2000). wingless and its signalling pathway have common and separable functions during tracheal development. Development 127, 4407–4417.

Llimargas, M. and Casanova, J. (1999). EGF signalling regulates cell invagination as well as cell migration during formation of tracheal system in Drosophila. Dev Gene Evol 209, 174–179.

Loganathan, R., Little, C. D. and Rongish, B. J. (2020). Extracellular matrix dynamics in tubulogenesis. Cellular Signalling 72, 109619.

Manning, G. and Krasnow, M. A. (1993). Development of the Drosophila tracheal system. In The development of Drosophila melanogaster., pp. 609–685. New York: Cold Spring Harbor Press.

Martin, D., Zusman, S., Li, X., Williams, E. L., Khare, N., DaRocha, S., Chiquet-Ehrismann, R. and Baumgartner, S. (1999). wing blister, A New Drosophila Laminin α Chain Required for Cell Adhesion and Migration during Embryonic and Imaginal Development. J Cell Biol 145, 191–201.

Olivares-Castiñeira, I. and Llimargas, M. (2017). EGFR controls Drosophila tracheal tube elongation by intracellular trafficking regulation. PLOS Genetics 13, e1006882.

Öztürk-Çolak, A., Marygold, S. J., Antonazzo, G., Attrill, H., Goutte-Gattat, D., Jenkins, V. K., Matthews, B. B., Millburn, G., Dos Santos, G., Tabone, C. J., et al. (2024). FlyBase: updates to the *Drosophila* genes and genomes database. GENETICS 227, iyad211.

Page-McCaw, A. and Ferrell, N. (2025). Basement membrane structure and function: Relating biology to mechanics. Matrix Biology 141, 16–31.

Paulissen, G., El Hour, M., Rocks, N., Guéders, M. M., Bureau, F., Foidart, J.-M., Lopez-Otin, C., Noel, A. and Cataldo, D. D. (2012). Control of Allergen-Induced Inflammation and Hyperresponsiveness by the Metalloproteinase ADAMTS-12. J Immunol 189, 4135–4143.

Pham, M. N., Schuweiler, M. and Ismat, A. (2018). The extracellular protease AdamTS-B inhibits vein formation in the Drosophila wing. genesis 56, e23255.

Port, F. and Bullock, S. L. (2016). Augmenting CRISPR applications in Drosophila with tRNA-flanked sgRNAs. Nat Methods 13, 852–854.

R Core Team (2021). R: A language and environment for statistical computing. R Foundation for Statistical Computing.

Rajković, A., Beracochea, M., Rogers, A. B., Eddy, S. R., Carter, N. P. and Finn, R. D. (2026). HMMER web server: 2026 update. Nucleic Acids Research 54, W272–W278.

Rocks, N., Paulissen, G., El Hour, M., Quesada, F., Crahay, C., Gueders, M., Foidart, J. M., Noel, A. and Cataldo, D. (2008). Emerging roles of ADAM and ADAMTS metalloproteinases in cancer. Biochimie 90, 369–379.

Sánchez-Cisneros, L. E., Frutis-Osorio, M. F. and Ríos-Barrera, L. D. (2025). A tale of two tissues: Patterning of the epidermis through morphogens and their role in establishing tracheal system organization. Cells & Development 184, 203998.

Schindelin, J., Arganda-Carreras, I., Frise, E., Kaynig, V., Longair, M., Pietzsch, T., Preibisch, S., Rueden, C., Saalfeld, S., Schmid, B., et al. (2012). Fiji: an open-source platform for biological-image analysis. Nat Methods 9, 676–682.

Scholl, A., Ndoja, I. and Jiang, L. (2021). Drosophila Trachea as a Novel Model of COPD. International Journal of Molecular Sciences 22, 12730.

Schulze, J. and Töpfer, U. (2025). Role of AdamTS-B in Drosophila wing vein formation. microPublication Biology.

Sherwood, D. R. (2021). Basement membrane remodeling guides cell migration and cell morphogenesis during development. Current Opinion in Cell Biology 72, 19–27.

Stanton, H., Melrose, J., Little, C. B. and Fosang, A. J. (2011). Proteoglycan degradation by the ADAMTS family of proteinases. Biochimica et Biophysica Acta (BBA) - Molecular Basis of Disease 1812, 1616–1629.

Taye, N., Singh, M., Baldock, C. and Hubmacher, D. (2023). Secreted ADAMTS-like 2 promotes myoblast differentiation by potentiating WNT signaling. Matrix Biology 120, 24–42.

Thuringer, A., Steinmetz, E., Abdullahi, M. and Ismat, A. (2026). The extracellular protease AdamTS-B is a dosage-sensitive regulator of unicellular tracheal branch morphogenesis. Biol Open 15, bio062602.

Töpfer, U. (2023). Basement membrane dynamics and mechanics in tissue morphogenesis. Biology open 12, bio059980.

Töpfer, U., Bischoff, M. C., Bartkuhn, M. and Holz, A. (2019). Serpent/dGATAb regulates Laminin B1 and Laminin B2 expression during Drosophila embryogenesis. Sci Rep 9, 15910.

Töpfer, U., Ryu, J., Guerra Santillán, K. Y., Schulze, J., Fischer-Friedrich, E., Tanentzapf, G. and Dahmann, C. (2024). AdamTS proteases control basement membrane heterogeneity and organ shape in Drosophila. Cell Reports 43, 114399.

Tsarouhas, V., Senti, K.-A., Jayaram, S. A., Tiklová, K., Hemphälä, J., Adler, J. and Samakovlis, C. (2007). Sequential Pulses of Apical Epithelial Secretion and Endocytosis Drive Airway Maturation in Drosophila. Developmental Cell 13, 214–225.

Urbano, J. M., Torgler, C. N., Molnar, C., Tepass, U., López-Varea, A., Brown, N. H., de Celis, J. F. and Martín-Bermudo, M. D. (2009). Drosophila laminins act as key regulators of basement membrane assembly and morphogenesis. Development 136, 4165–4176.

Wolfstetter, G. and Holz, A. (2012). The role of LamininB2 (LanB2) during mesoderm differentiation in Drosophila. Cell. Mol. Life Sci. 69, 267–282.

Wolfstetter, G., Dahlitz, I., Pfeifer, K., Töpfer, U., Alt, J. A., Pfeifer, D. C., Lakes-Harlan, R., Baumgartner, S., Palmer, R. H. and Holz, A. (2019). Characterization of Drosophila Nidogen/entactin reveals roles in basement membrane stability, barrier function and nervous system patterning. Development 146, dev168948.

Yarnitzky, T. and Volk, T. (1995). Laminin Is Required for Heart, Somatic Muscles, and Gut Development in the *Drosophila* Embryo. Developmental Biology 169, 609–618.

Yurchenco, P. D. (2011). Basement Membranes: Cell Scaffoldings and Signaling Platforms. Cold Spring Harbor Perspectives in Biology 3, a004911–a004911.

